# Association of spinocerebellar ataxia related variants with myokymia and neuromyotonia in dogs

**DOI:** 10.1101/2022.11.17.516927

**Authors:** An Vanhaesebrouck, Mario Van Poucke, Kimberley Stee, Luc Peelman, Luc Van Ham, Sofie F.M. Bhatti

## Abstract

**Background:** KCNJ10 and CAPN1 variants have been shown to cause spinocerebellar ataxia (SCA) in Jack Russell, Parson Russell and Fox terriers (JRT, PRT and FT). However, their association with the clinical manifestation of myokymia and neuromyotonia (M/NM), often reported alongside SCA, remains unclear.

**Aims:** To investigate the association between M/NM and SCA-related variants in 34 with SCA and/or M/NM affected dogs (30 JRTs; 1 PRT; 1 Yorkshire terrier, YT; 1 Dachshund; 1 crossbreed).

**Methods:** KCNJ10 XM_038448705.1:c.627C>G (p.(Ile209Met)) and CAPN1 XM_038425033.1:c.344G>A (p.(Cys115Tyr)) variants, and the complete coding sequence (CDS) of KCNA1, KCNA2, KCNA6, KCNJ10 and HINT1, were analysed via Sanger sequencing.

**Results:** The KCNJ10 c.627C>G variant was homozygously present in 16 JRTs, 1 Dachshund and 1 crossbreed with SCA and M/NM, and in 9 JRTs with SCA but without M/NM. The CAPN1 c.344G>A variant was homozygously present in 1 PRT with SCA but without M/NM. Both variants were not found in 2 JRTs with SCA but without M/NM, neither in 3 JRTs and 1 YT without SCA but with M/NM. No other causal variants were found in the coding sequence of the investigated candidate genes in these latter 6 dogs.

**Conclusions:** The KCNJ10 c.627C>G or CAPN1 c.344G>A variant was confirmed to be the causal variant in 28 of the 34 affected dogs (including 1 Dachshund with the KCNJ10 variant). The fact that 10 of these 28 dogs did not suffer from M/NM (all of them suffered from SCA) and that these variants were not found in 4 dogs suffering from M/NM without SCA, suggests that M/NM is caused by another genetic variant or that these patients suffer from an acquired (immune-mediated) syndrome, similar to humans. Another SCA-causing variant is probably also segregating in JRTs (but not in the CDS of the investigated candidate genes), because no causal variant was found in 2 JRTs suffering from SCA without M/NM.

## INTRODUCTION

Generalized myokymia and neuromyotonia (M/NM) are clinical manifestations of hyperactivity of the peripheral motor nerves. Myokymia is characterized by vermicular muscle movements while neuromyotonia is characterized by generalized, severe and long lasting muscle stiffness (Vanhaesebrouck et al., 2013). Generalized M/NM are mainly observed in humans as an immune-mediated disorder, in which the immune system directs itself to proteins related to the potassium channels of the peripheral nerve (KCNA1, KCNA2 and KCNA6; Newsom-Davis, 1997). It has also been described as a hereditary disorder in children related to variants in certain potassium channels of the peripheral nerve (KCNA1 and KCNQ2; Dedek et al., 2001; Poujois et al., 2006). Variants in HINT1 have shown to cause neuromyotonia and axonal neuropathy (Zimoń et al., 2012). Lastly, peripheral hyperexcitability can be caused by electrolyte disturbances of calcium and magnesium (Wijnberg et al., 2002).

In Jack and Parson Russell terriers (JRT and PRT) a syndrome characterized by a typical dancing bouncing gait has been described for the first time in the seventies, and at that point the disease was termed “hereditary ataxia” (Hartley and Palmer, 1973). Histopathological features are characterized by degenerative axonal changes and demyelinisation within the spinal cord, brainstem and nerve roots. Since 2004, several case series described that JRTs with “hereditary ataxia” often also suffered from M/NM (Van Ham et al., 2004; Vanhaesebrouck et al., 2010; Bhatti et al., 2011). In 2014, a variant in KCNJ10 was linked to JRTs and PRTs with hereditary spinocerebellar ataxia (SCA), with or without M/NM and seizures (Gilliam et al., 2014). This variant was also found in Fox terriers (FT; Rohdinetal., 2015). In addition, a variant in CAPN1 has been found in PRTs with hereditary late-onset ataxia (LOA; Forman et al., 2013). LOA has so far not been described with clinical signs of M/NM.

In humans, KCNJ10 variants have been linked to Epilepsy Ataxia Sensorineural Deafness Tubulopathy (EAST syndrome; Bockenhauer et al., 2009). Neither myokymia nor neuromyotonia has been reported in KCNJ10 variants in humans or in experimental animal models (Neusch et al., 2001). In addition, no electrodiagnostic tests have been reported in this syndrome in humans. Little is known of CAPN1 variants in other species than dogs.

This study aims to further clarify the association between clinical features of M/NM and SCA-related variants in 34 affected dogs (30 JRTs; 1 PRT; 1 Yorkshire terrier, YT; 1 Dachshund; 1 crossbreed).

## METHODS

### Cases

Thirty-three dogs (30 JRTs, 1 PRT, 1 Yorkshire terrier (YT) and 1 Dachshund) suffering from SCA and/or M/NM with a time of onset less than 3 years, were clinically described in detail in previous studies, without association of their clinical presentation with a genetic variant (Van Ham et al., 2004; Vanhaesebrouck et al., 2010; Bhatti et al., 2011; Vanhaesebrouck et al., 2011). SCA was defined as a dancing bouncing gait, involving all limbs but worse on the hindlimbs, and including a head tremor. Myokymia was defined as vermicular movements of the skin or mucosa overlying the muscle, present in more than 2 muscles. Neuromyotonia was defined as a long lasting generalized stiffness with the dog in lateral decubitus with rigid limbs for more than 10 minutes, associated with hyperthermia and without any loss of consciousness (Vanhaesebrouck et al., 2013). A crossbreed suffering from SCA with M/NM with a time of onset less than 3 years and tested for both KCNJ10 c.627C>G and CAPN1 c.344G>A variants (Alexandria Haimbaugh, pers. comm.) was added in this paper (Table 1).

**Table 1.** Information on the breed, phenotype (SCA and M/NM) and genotype (KCNJ10 and CAPN1) of the 34 investigated dogs.

| Breed | SCA | M/NM | KCNJ10 | CAPN1 | Number |
| --- | --- | --- | --- | --- | --- |
| JRT | + | + | **G/G** | G/G | 16 |
| Dachshund | + | + | **G/G** | G/G | 1 |
| Mixed-breed | + | + | **G/G** | G/G | 1 |
| JRT | + | - | **G/G** | G/G | 9 |
| PRT | + | - | *C/C* | **A/A** | 1 |
| JRT | + | - | *C/C* | G/G | 2 |
| JRT | - | + | *C/C* | G/G | 3 |
| YT | - | + | *C/C* | G/G | 1 |

CAPN1, XM\_038425033.1:c.344G>A (p.(Cys115Tyr)) CAPN1 genotype; JRT, Jack Russell terrier; KCNJ10, XM\_038448705.1:c.627C>G (p.(Ile209Met)) KCNJ10 genotype; M/NM, myokymia and/or neuromyotonia; PRT, Parson Russell terrier; SCA, spinocerebellar ataxia; YT, Yorkshire terrier. Variant alleles are in bold.

### Genetic analysis

Genomic DNA was isolated from EDTA-blood samples. Sanger sequencing was used to genotype the KCNJ10 XM_038448705.1:c.627C>G (p.(Ile209Met)) variant (Gilliam et al., 2014) and the CAPN1 XM_038425033.1:c.344G>A (p.(Cys115Tyr)) variant (Forman et al., 2013). Dogs in which the phenotype could not be explained by any of the 2 tested variants, were screened for causal variants in the coding sequence (CDS) of candidate genes KCNA1, KCNA2, KCNA6, KCNJ10 and HINT1 via Sanger sequencing. Details of the genetic analysis is given in Table 2.

**Table 2.**
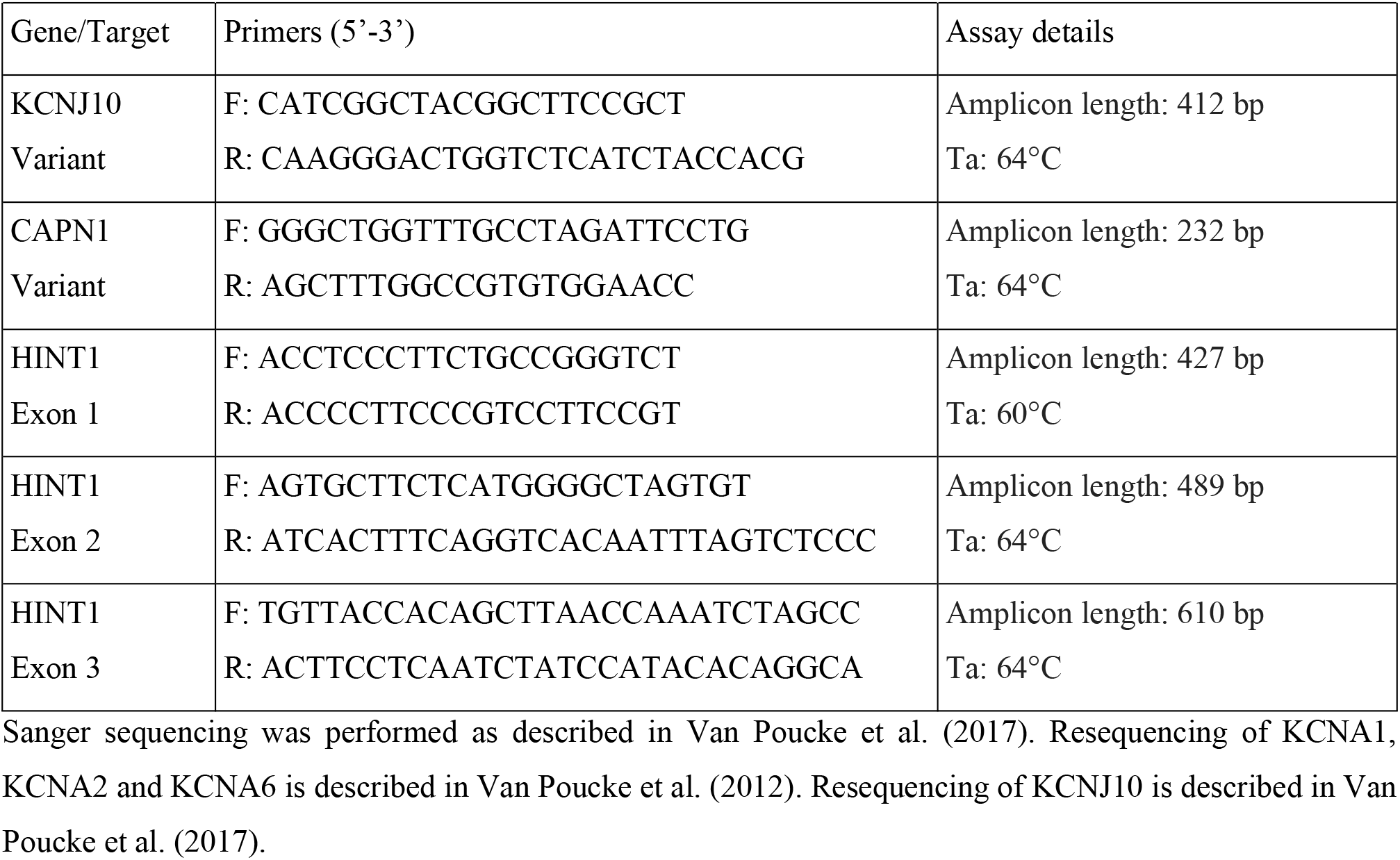
Details on the genetic analysis.

## RESULTS

All 18 dogs suffering from SCA with M/NM (16 JRTs, 1 Dachshund and 1 crossbreed), were homozygous for the KCNJ10 c.627C>G variant. Also 9 of the 11 JRTs suffering from SCA but without M/NM were homozygous for this KCNJ10 variant, while the other 2 did not carry the variant. The only PRT, also suffering from SCA but without M/NM, did not carry this KCNJ10 variant, but was homozygous for the CAPN1 c.344G>A variant. All 4 dogs suffering from M/NM but without SCA did not carry any of the 2 variants (Table 1). CDS analysis of KCNA1, KCNA2, KCNA6, KCNJ10 and HINT1 in the 6 dogs in which the phenotype could not be explained by the above mentioned KCNJ10 and CAPN1 variants, did not reveal any other causal variant.

## DISCUSSION

The KCNJ10 c.627C>G variant was confirmed to be the causal variant in 27 affected dogs. The variant, already described in JRTs, PRTs and FTs (Gilliam et al., 2014; Rohdin et al., 2015), was not only found in 25 JRTs but also in a crossbreed and a Dachshund. Based on the crossbreed’s physical appearance, JRT ancestry seems likely, which would explain the origin of the variant. In the Dachshund, it is the first report of the presence of this variant. Although the exact origin of the Dachshund breed remains unclear, they were most likely crossed with terriers in the past (Loeffler, 1891), which would explain the presence of this variant in this breed. The CAPN1 c.344G>A variant, described in PRTs (Forman et al., 2013), was confirmed to be the causal variant in 1 affected PRT. The fact that 10 of these 28 affected dogs with a known causal variant, did not suffer from M/NM (all of them suffered from SCA) and that these known causal variants were not found in 4 dogs (3 JRTs and 1 YT) presenting classic M/NM but lacking SCA (a manifestation previously also described by Reading and Mckerrell, 1993), suggests that M/NM is caused by another genetic variant or that these patients suffer from an acquired (immune-mediated) syndrome, similar to humans (Irani et al., 2012).

For 2 JRTs with SCA but without M/NM, no genetic variant was found. This suggest that another causal variant is segregating in the JRT breed, but not in the CDS of KCNA1, KCNA2, KCNA6, KCNJ10 and HINT1.

## ABBREVIATIONS

CAPN1: calpain 1
CDS: coding sequence
EAST: Epilepsy Ataxia Sensorineural Deafness Tubulopathy
FT: Fox terrier
HINT1: histidine triad nucleotide binding protein 1
JRT: Jack Russell terrier
KCNAx: potassium voltage-gated channel subfamily A member x
KCNJ10: potassium inwardly rectifying channel subfamily J member 10
LOA: late onset ataxia
M/NM: myokymia and neuromyotonia
PCR-RFLP: polymerase chain reaction – restriction length polymorphism
PRT: Parson Russell terrier
SCA: spinocerebellar ataxia
YT: Yorkshire terrier

## ACKNOWLEDGEMENTS

We would like to thank Dominique Vander Donckt, Linda Impe, Carolien Rogiers and Ruben Van Gansbeke for assistance in the genetic analysis of the dogs. We are grateful for the help of the breeders and the owners of the affected dogs.

## REFERENCES

Bhatti SF, Vanhaesebrouck AE, Van Soens I, Martlé VA, Polis IE, Rusbridge C, Van Ham LM. Myokymia and neuromyotonia in 37 Jack Russell terriers. Vet J. 2011 Sep;189(3):284–8. doi: 10.1016/j.tvjl.2010.07.011.

Bockenhauer D, Feather S, Stanescu HC, Bandulik S, Zdebik AA, Reichold M, Tobin J, Lieberer E, Sterner C, Landoure G, Arora R, Sirimanna T, Thompson D, Cross JH, van’t Hoff W, Al Masri O, Tullus K, Yeung S, Anikster Y, Klootwijk E, Hubank M, Dillon MJ, Heitzmann D, Arcos-Burgos M, Knepper MA, Dobbie A, Gahl WA, Warth R, Sheridan E, Kleta R. Epilepsy, ataxia, sensorineural deafness, tubulopathy, and KCNJ10 mutations. N Engl J Med. 2009 May 7;360(19):1960–70. doi: 10.1056/NEJMoa0810276.

Dedek K, Kunath B, Kananura C, Reuner U, Jentsch TJ, Steinlein OK. Myokymia and neonatal epilepsy caused by a mutation in the voltage sensor of the KCNQ2 K+ channel. Proc Natl Acad Sci USA. 2001 Oct 9;98(21):12272–7. doi: 10.1073/pnas.211431298.

Forman OP, De Risio L, Mellersh CS. Missense mutation in CAPN1 is associated with spinocerebellar ataxia in the Parson Russell Terrier dog breed. PLoS One. 2013 May 31;8(5):e64627. doi: 10.1371/journal.pone.0064627.

Gilliam D, O’Brien DP, Coates JR, Johnson GS, Johnson GC, Mhlanga-Mutangadura T, Hansen L, Taylor JF, Schnabel RD. A homozygous KCNJ10 mutation in Jack Russell Terriers and related breeds with spinocerebellar ataxia with myokymia, seizures, or both. J Vet Intern Med. 2014 May-Jun;28(3):871–7. doi: 10.1111/jvim.12355.

Hartley WJ, Palmer AC. Ataxia in Jack Russell terriers. Acta Neuropathol. 1973;26(1):71–4. doi: 10.1007/BF00685524.

Irani SR, Pettingill P, Kleopa KA, Schiza N, Waters P, Mazia C, Zuliani L, Watanabe O, Lang B, Buckley C, Vincent A. Morvan syndrome: clinical and serological observations in 29 cases. Ann Neurol. 2012 Aug;72(2):241–55. doi: 10.1002/ana.23577.

Loeffler W. Dachshunds. In: Go Shields, ed. The American Book of the Dog. University of California: Cassell and Co; 1891:217–239. doi: 10.5962/bhl.title.20049.

Neusch C, Rozengurt N, Jacobs RE, Lester HA, Kofuji P. Kir4.1 potassium channel subunit is crucial for oligodendrocyte development and in vivo myelination. J Neurosci. 2001 Aug 1;21(15):5429–38. doi: 10.1523/JNEUROSCI.21-15-05429.2001.

Newsom-Davis J. Autoimmune neuromyotonia (Isaacs’ syndrome): an antibody-mediated potassium channelopathy. Ann NY Acad Sci. 1997 Dec 19;835:111–9. doi: 10.1111/j.1749-6632.1997.tb48622.x.

Poujois A, Antoine JC, Combes A, Touraine RL. Chronic neuromyotonia as a phenotypic variation associated with a new mutation in the KCNA1 gene. J Neurol. 2006 Jul;253(7):957–9. doi: 10.1007/s00415-006-0134-y.

Reading MJ, McKerrell RE. Suspected myokymia in a Yorkshire terrier. Vet Rec. 1993 Jun 5;132(23):587–8. doi: 10.1136/vr.132.23.587.

Rohdin C, Gilliam D, O’Leary CA, O’Brien DP, Coates JR, Johnson GS, Jäderlund KH. A KCNJ10 mutation previously identified in the Russell group of terriers also occurs in Smooth-Haired Fox Terriers with hereditary ataxia and in related breeds. Acta Vet Scand. 2015 May 23;57(1):26. doi: 10.1186/s13028-015-0115-1.

Vanhaesebrouck AE, Van Soens I, Poncelet L, Duchateau L, Bhatti S, Polis I, Diels S, Van Ham L. Clinical and electrophysiological characterization of myokymia and neuromyotonia in Jack Russell Terriers. J Vet Intern Med. 2010 Jul-Aug;24(4):882–9. doi: 10.1111/j.1939-1676.2010.0525.x.

Vanhaesebrouck AE, Bhatti SF, Polis IE, Plessas IN, Van Ham LM. Neuromyotonia in a dachshund with clinical and electrophysiological signs of spinocerebellar ataxia. J Small Anim Pract. 2011 Oct;52(10):547–50. doi: 10.1111/j.1748-5827.2011.01123.x.

Vanhaesebrouck AE, Bhatti SF, Franklin RJ, Van Ham L. Myokymia and neuromyotonia in veterinary medicine: a comparison with peripheral nerve hyperexcitability syndrome in humans. Vet J. 2013 Aug;197(2):153–62. doi: 10.1016/j.tvjl.2013.03.002.

Van Ham L, Bhatti S, Polis I, Fatzer R, Braund K, Thoonen H. ‘Continuous muscle fibre activity’ in six dogs with episodic myokymia, stiffness and collapse. Vet Rec. 2004 Dec 11;155(24):769–74. doi: 10.1136/vr.155.24.769.

Van Poucke M, Vanhaesebrouck AE, Peelman LJ, Van Ham L. Experimental validation of in silico predicted KCNA1, KCNA2, KCNA6 and KCNQ2 genes for association studies of peripheral nerve hyperexcitability syndrome in Jack Russell Terriers. Neuromuscul Disord. 2012 Jun;22(6):558–65. doi: 10.1016/j.nmd.2012.01.008.

Van Poucke M, Stee K, Bhatti SF, Vanhaesebrouck A, Bosseler L, Peelman LJ, Van Ham L. The novel homozygous KCNJ10 c.986T>C (p.(Leu329Pro)) variant is pathogenic for the SeSAME/EAST homologue in Malinois dogs. Eur J Hum Genet. 2017 Feb;25(2):222–226. doi: 10.1038/ejhg.2016.157.

Wijnberg ID, van der Kolk JH, Franssen H, Breukink HJ. Electromyographic changes of motor unit activity in horses with induced hypocalcemia and hypomagnesemia. Am J Vet Res. 2002 Jun;63(6):849–56. doi: 10.2460/ajvr.2002.63.849.

Zimoń M, Baets J, Almeida-Souza L, De Vriendt E, Nikodinovic J, Parman Y, Battaloğlu E, Matur Z, Guergueltcheva V, Tournev I, Auer-Grumbach M, De Rijk P, Petersen BS, Müller T, Fransen E, Van Damme P, Löscher WN, Barišić N, Mitrovic Z, Previtali SC, Topaloğlu H, Bernert G, Beleza-Meireles A, Todorovic S, Savic-Pavicevic D, Ishpekova B, Lechner S, Peeters K, Ooms T, Hahn AF, Züchner S, Timmerman V, Van Dijck P, Rasic VM, Janecke AR, De Jonghe P, Jordanova A. Loss-of-function mutations in HINT1 cause axonal neuropathy with neuromyotonia. Nat Genet. 2012 Oct;44(10):1080–3. doi: 10.1038/ng.2406.

